# Imaging real-time tactile interaction with two-person dual-coil fMRI

**DOI:** 10.1101/861252

**Authors:** Ville Renvall, Jaakko Kauramäki, Sanna Malinen, Riitta Hari, Lauri Nummenmaa

## Abstract

Studies of brain mechanisms supporting social interaction are demanding because real interaction only occurs when the persons are in contact. Instead, most brain imaging studies scan subjects individually. Here we present a proof-of-concept demonstration of two-person blood oxygenation dependent (BOLD) imaging of brain activity from two individuals interacting inside the bore of a single MRI scanner. We developed a custom 16-channel (8 + 8 channels) two-helmet coil with two separate receiving elements providing whole-brain coverage, while bringing participants into a shared physical space and realistic face-to-face contact. Ten subject pairs were scanned with the setup. During the experiment, subjects took turns in tapping each other’s’ lip versus observing and feeling the taps timed by auditory instructions. Networks of sensorimotor brain areas were engaged alternatingly in the subjects during executing motor actions as well as observing and feeling them; these responses were clearly distinguishable from the auditory responses occurring similarly in both participants. Even though the signal-to-noise ratio of our coil system was compromised compared with standard 32-channel head coils, our results show that the two-person fMRI scanning is a feasible in studying the brain basis of social interaction.

## Introduction

Humans are embedded in complex social networks where individuals interact at different temporal scales. Most social interactions, such as verbal and nonverbal communications, occur in dyads or groups, where people constantly strive to predict, understand and influence each other. During the interaction, sensory, cognitive and emotional information is constantly remapped in the observers’ brain and used for motor actions as responses attuned to the received input (Hari & Kujala, 2009). Thus the interlocutors’ minds are intertwined into a shared system facilitating reciprocation (Hasson et al., 2012; Nummenmaa et al., 2018; Smirnov et al., 2017) as well as anticipation of the other person’s acts, allowing distribution of neural processing across brains to aid, for example, problem solving.

Some aspects of human social behaviour – in particular perceptual and decision-making processes - can be studied by measuring single brains in isolation. Conventional BOLD-fMRI experiments allow localization of different social processes in the brain, while intersubject correlation (ISC) analysis based on voxelwise temporal correlation of BOLD-fMRI time series (Bartels & Zeki, 2004; Hasson et al., 2004; Wilson et al., 2008) or neuromagnetic activity with higher temporal resolution (Hari et al., 2015) across subject pairs can be used to index similarity of sensory and socioemotional information processing across subjects (Lahnakoski et al., 2014; Nummenmaa et al., 2012), but also similarity of person preferences and social relationships (Parkinson et al., 2018). Although this kind of data-driven analyses can be used to tackle questions such as social perception with high-dimensional stimulus spaces, they are still essentially based on measurement of extrinsic, fixed stimuli and lack the very definition of social interaction, as the subjects have no influence whatsoever on other peoples’ minds in the experiment. This is a critical limitation as it has been argued that social interaction cannot be reduced to sequential, partially parallel processing of the input of the interacting brains, because social interaction only emerges when the two brains (via their owners) are hooked up together (Hari et al., 2015; Hari et al., 2016; Nummenmaa et al., 2018). Simply put, real-time social interaction does not exist when two or more individuals are not engaged in the same physical or virtual space (De Jaegher et al., 2010).

Against this background, reciprocal social cognitive processes cannot be understood completely without studying the complete interaction unit consisting of two individuals (Konvalinka & Roepstorff, 2012). Behavioral work suggests that social interaction tunes the individuals into a self-organizing, interactive state. For example, humans automatically mimic others emotional expressions (Dimberg et al., 2000), gaze direction (Nummenmaa & Calder, 2009) and postures (Lakin et al., 2003). Social signals, such as laughter, also automatically tune individuals in not just at the level of motor responses, but also in terms of activation of specific neurotransmitter systems (Manninen et al., 2017). Moreover, many social processes, such as gaze following (Frischen et al., 2007; Pfeiffer et al., 2013) and turn taking during conversation (Stivers et al., 2009), takes place with gaps less than 250 ms, and social interaction may lead to episodes of two-person flow without neither of them consciously leading or following (Noy et al., 2011). Yet, most of what we know about human social brain functions comes from “spectator” studies where the brains are assumed to generate responses to pre-defined stimulation (Hari et al., 2015). Even though this approach has been successful in delineating the brain basis of social perception, and on some occasions of social communication, it tells relatively little about the actual mechanisms of dynamic social interaction. Consequently, several researchers have suggested that the spectator paradigm and offline social cognition studies should be complemented with real-time two-person paradigms, where two interacting individuals constantly generate “stimuli” for each other (Hari & Kujala, 2009; Hasson et al., 2012; Nummenmaa et al., 2018; Redcay & Schilbach, 2019; Schilbach et al., 2013).

Still, some aspects of human communication can be investigated using alternated scanning of the subjects sending and receiving information. In such an approach, the senders convey some social information via, for example, speech or gestures, while their brain activity as well as the communicative information are recorded. The communicative information can then be presented to the receiver subjects as stimuli during brain imaging, allowing joint analysis of the brain activity of the sender and receiver subjects. This line of work has revealed how successful communication via speech (Smirnov et al., 2019; Stephens et al., 2010), hand gestures (Schippers et al., 2010; Smirnov et al., 2017) and facial expressions (Anders et al., 2011) enhances similarity of neural activation patterns across the interlocutors in a task-specific manner. This approach however lacks any interactivity, as the receiver subjects are essentially viewing pre-recorded stimuli, and need not to generate any responses to them. Recently different neuroimaging techniques have been proposed for studying dynamic “live” interaction. In a hyperscanning approach, two individuals are scanned with two MRI (King-Casas et al., 2008; Montague et al., 2002; Saito et al., 2010), MEG (Baess et al., 2012), devices connected with an audio-video link, thus enabling interaction of two subjects in independent devices. Furthermore, with dyadic EEG recordings real face-to-face to interaction can be achieved reasonably unconstrained social interaction tasks (Babiloni & Astolfi, 2014). The resulting natural sense of presence of another individual might be critical for understanding the brain basis of social interaction. For example, resting-state brain activity in nonhuman primates is different when conspecifics are present versus absent (Monfardini et al., 2016). In humans, interaction with real rather than recorded persons elicits stronger heamodynamic responses (Redcay et al., 2010), and even early electrophysiological responses such as the face-sensitive N170 responses are amplified for real human faces versus those of a human-like dummy (Ponkanen et al., 2008). These findings highlight the importance of co-presence with other people, and the consequent changes in the way the brain processes both internal and external cues. Consequently, to understand the intricacies of the brain basis of human social interaction and communication, we need techniques that allow simultaneous recording of from two individuals in the same physical space as has already been technically achieved with simultaneous EEG (e.g. Dumas et al., 2010) and NIRS (e.g. Cui et al., 2012; Funane et al., 2011) recordings, as these devices can be easily attached to subjects measured in a conventional face-to-face settings. Nevertheless, neither of these techniques allow volumometric measurements of the deep brain structures, many of which are critical for human social processes (Lahnakoski et al., 2012; Saarimaki et al., 2018; Saarimäki et al., 2016).

## The current study

One potentially powerful approach for studying brain basis of social interaction involves simultaneous blood oxygenation dependent (BOLD) imaging of two persons within one magnetic resonance imaging scanner. Such an approach would bring both subjects into the same physical space whilst allowing tomographic imaging of haemodynamic brain activation. Currently, one such solution has been published, based on decoupled circular-polarized volume coil for two heads (Lee, 2015; Lee et al., 2012). We have, in turn, developed a custom-built 16-channel (8 + 8 channels) two-helmet coil with two separate receiving elements (Renvall & Malinen, 2012, OHBM Conference), resulting in experimental setups where the subjects were facing each other so that their feet pointed to opposite directions in the magnet bore. In the present proof-of-concept study demonstrate how haememodynamic signals can be recorded during real-time social interaction using this a novel MRI receive coil modified so that the subject can lie parallel to each other while sharing the same physical space in a realistic face-to-face contact. The setup thereby allows seamless interaction between the members of the dyad, while providing whole-brain coverage.

Because this was the very first proof-of-concept human experiment done with the dual coil, we wanted to benchmark the feasibility of the setup with a robust and simple social interaction task, rather than setting up an overly complex design without knowing the potential limitations of the coil setup. Consequently, we used social touching as the model task, as touching is an intimate way of conveying affect and trust in social relationships (Nummenmaa et al., 2016; Suvilehto et al., 2015; Suvilehto et al., 2019). Because relationship-social touching cannot be adequately simulated by experimental tools or apparatuses, its investigation would benefit from the real-time dyadic imaging. During the fMRI experiments, subjects took turns in tapping each others’ lip versus observing and feeling the taps. We show that overlapping networks of sensorimotor brain areas are engaged during executing motor actions as well as observing and feeling them, suggesting that the two-person fMRI recordings are feasible for studying the brain basis of social interaction.

## Materials and Methods

### Subjects

We scanned 10 pairs of volunteers with a mean age of and 23 ± 3 years (20 subjects; 7 female–male pairs and 3 female–female pairs). One further pair was scanned but excluded due to excessive head motions: one of the subjects moved so that the detector array’s sensitivity was compromised and repositioning was not possible due to time constraints. All subjects were right handed per self-report, and none of them reported any history of neurological illness. All pairs were friends or romantic partners. The study was approved by the Aalto University Institutional Review Board. All subjects gave written informed consent and were screened for MRI exclusion criteria prior to scanning.

### MRI acquisition

Data were acquired with 3-T whole body MRI system (MAGNETOM Skyra 3.0 T, Siemens Healthcare, Erlangen, Germany) with both a vendor-provided 32-channel receive head coil (reference scans) and a custom-built 16-channel (8 + 8 channels) two-head, two-helmet receive coil (anatomical images, task-based fMRI, and resting state scan). With both receive coils, the integrated body coil was used for transmit. **Figure 1** shows an overview of the coil and subject setup. As already mentioned, we originally experimented with a setup where subjects were lying either sideways or in a supine position, while entering the gantry from the opposite ends so that a second custom MRI bed was used for the backwards entry. This setup was however discarded due to subject discomfort and concomitant motion-related artifacts.

**Figure 1.**
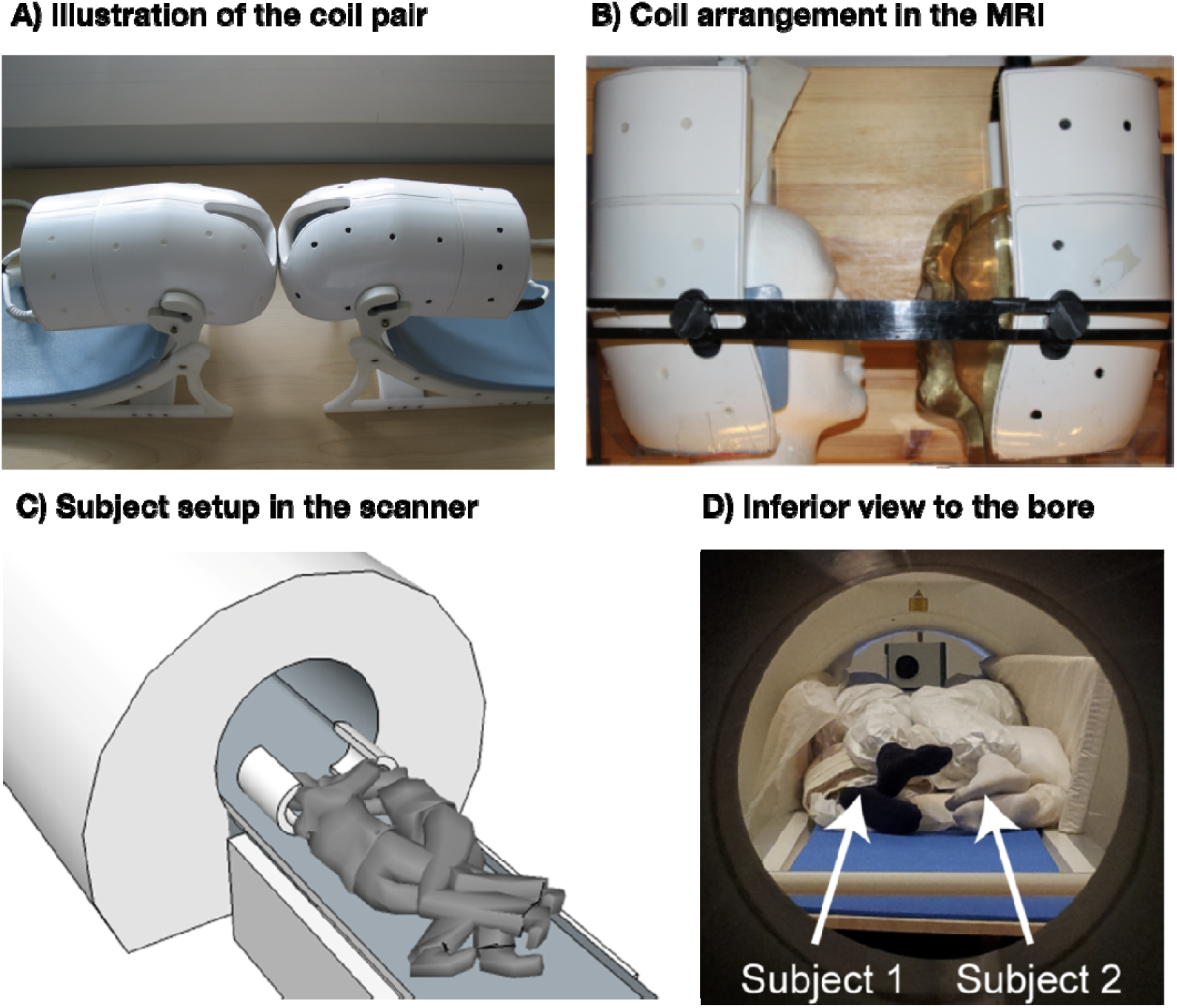
Coil and subject setup. (A–B) Illustration of the dual coil and its arrangement in the scanner. (C–D) Subject setup inside the scanner.

Every scanning session consisted of two parts. First both subjects were scanned one-by-one using normal one-person setup (head-first supine, 32-channel coil). T1-weighted MP-RAGE images were acquired for anatomical reference, and gradient echo (GRE) echo-planar imaging (EPI) data were acquired for evaluating the temporal signal-to-noise ratio (tSNR), especially in comparison with the two-person data. Imaging parameters for the MP-RAGE scans were as follows: repetition time (TR) = 2.53 s, echo time (TE) = 28 ms, readout flip angle (*α*) = 7°, 256 × 256 × 176 imaging matrix, isotropic 1-mm^3^ resolution, and GRAPPA reduction factor (*R*) = 2. The parameters for the GRE-EPI were: TR = 3 s, TE = 28 ms, *a* = 80°, fat saturation was used, in-plane imaging matrix (frequency encoding × phase encoding) = 70 × 70, field-of-view (FOV) = 21 × 21 cm^2^, in-plane resolution 3 × 3 mm^2^, R = 2, effective echo spacing (esp) = 0.26 ms, bandwidth = 2380 Hz/pixel (total bandwidth = 167 kHz), phase encoding in anterior–posterior direction, slice thickness = 3 mm with 10% slice gap, interleaved slice-acquisition order. Altogether 126 volumes, with 49 oblique axial slices in each, were acquired during the 6 min 18 s data-collection period. Three “dummy” scans were acquired at the beginning of each acquisition to stabilize the longitudinal magnetization.

Next the subjects were positioned in the scanner together with the two-head coil; the subjects were lying on their sides, facing each other at a close distance. Localizer and GRE-EPI data were acquired after shimming the magnet, using the semi-automated workflow by iteratively acquiring B_0_ field maps and calculating the shims for as long as the shim was deemed unacceptable. In this phase, the subjects could be repositioned if the their field maps appeared excessively dissimilar. The scan parameters were the same as in the one-person setup, with the following exceptions: in-plane matrix = 160 × 70, FOV = (48.6 × 20.1) cm^2^, and bandwidth = 2404 Hz/pixel (total bandwidth = 385 kHz). Moreover, the phase encoding was in subjects’ left–right direction to avoid aliasing ghosts from one subject’s brain into the other, and to reduce distortion and scan time by limiting the number of phase encoding steps (to 35 per slice). The 49 slices were oriented axially and tilted only to maximize the symmetry of the acquisition of the two brains. During the two-person measurements, the bodies of the subjects were in contact (without direct skin contact) and pillows and foam mattresses were used to make the subjects as comfortable as possible. The subjects’ heads were stabilized using small pillows with non-slippery surface and additional support was provided using a large vacuum pillow that once deflated retained its shape throughout the session.

### Touching task

**Figure 2** summarizes the structure of the tactile-motor task. The subjects took turns in repeatedly tapping (“actor” subject) the lower lip of their partner (“receiver” subject) with the tip of the index finger, so that both partners could also clearly see the finger movement. This site was chosen so that that the finger movements would be clearly visible to both subjects. Self-paced (~2 Hz) tapping was performed throughout the 30-s task blocks. Subjects were stressed to minimize finger movements, because motion near the imaging volume perturbs the magnetic field and can interfere with the spatial encoding and introduce head motion. During the rest blocks the subjects were instructed to hold their finger close by but not touching the lower lip of their partner. Each task run contained six rest–task block cycles with an additional rest block at the end of the run. Except for the initial rest condition, transitions between blocks were cued by pre-recorded voice command “Touch” and “Rest”. These were delivered to the participants by connecting the stimulus computer’s audio output to the magnet console to use the intercom system of the MRI scanner. Presentation software (Neurobehavioral Systems, Berkeley, CA, USA) controlled stimulus presentation. After the first task run we confirmed that the subjects could hear the voice commands. For any given run, only one of the subjects performed the active touching task while the other focused on feeling the taps. The roles were switched between runs. Both subjects were always scanned twice in both roles so that altogether 4 task runs with 126 EPI volumes in each were acquired.

**Figure 2.**
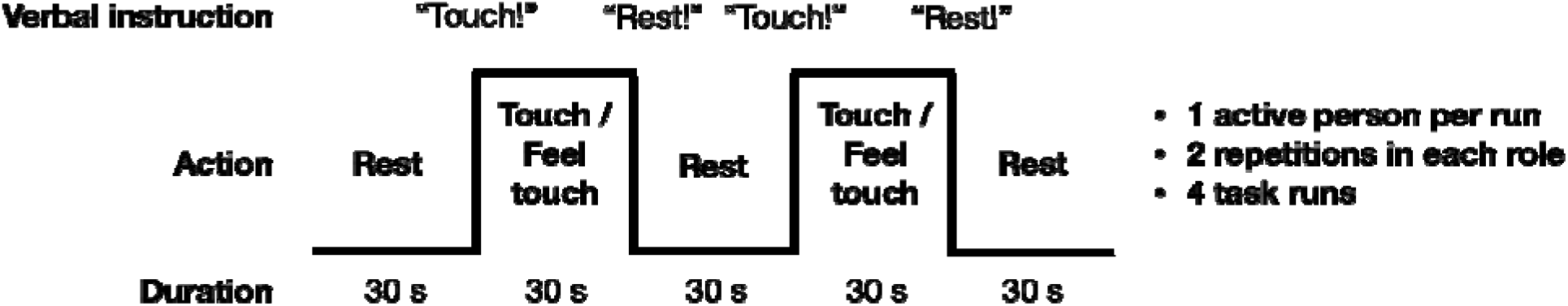
Experimental design. Subjects took 30-s turns in tapping the top of each other’s lip with their index finger, resulting in alternating tapping-feeling boxcar design with complete antiphase across the subjects. Turns were indicated with commands relayed via headphones.

### Resting-state scans

Resting state scans were obtained for inspecting signal quality. During the single-subject GRE-EPI data acquisition, the subjects were instructed to lie still with eyes open but not looking at each other. Eye-blinking was allowed. The two-person resting-state scans were always acquired prior to the task scans, asking the subjects to keep their eyes open and still.

### Image preprocessing – One-person scans

The fMRI data were preprocesses in Matlab utilizing custom code and FSL functions (Jenkinson et al., 2012). The one-person fMRI data were motion-corrected using FSL MCFLIRT (Jenkinson et al., 2002). Next, slice-timing correction was applied and the frame-wise motion within each fMRI run was corrected using FSL function MCFLIRT after brain extraction using BET (Smith, 2002) and smoothed with structure-preserving smoothing with SUSAN (Smith & Brady, 1997) that employed a 6-mm (full-width-at-half-maximum, FWHM) Gaussian smoothing kernel. The data were rigidly (6 free parameters) aligned to the anatomical MP-RAGE scans, with narrow search space for the alignment because the receiver intensity was spatially atypical and prohibited the use of the standard options for several datasets. The anatomical images were normalized to the MNI304 space, and the resulting warps were then applied to the EPI images. Data were finally smoothed using a Gaussian kernel with 8mm FWHM.

### Image preprocessing – Two-person scans

Individual heads were first separated and rotated to standard head-first supine orientation using a fixed coordinate transformation without resampling. Next both subjects’ data were preprocessed independently as described above. Preprocessing was concluded by recombining the data of each pair so that one subjects’ data were in MNI space, and the other subject’s data were placed nose-to-nose with that to mimic the actual positioning during the scanning.

### Signal-to-Noise Ratios

Coil performance was assessed with temporal signal-to-noise ratio (tSNR) of resting-state fMRI scans comprising of 126 time points. The FSL BET program was used to extract the brain voxels from the images, after which the data were motion-corrected using FSL MCFLIRT. Next, voxelwise tSNR values were calculated as the ratio of the mean signal over the measurement, divided by the standard deviation (std) at each voxel. For comparison, similar analysis was carried out for the one-person resting-state data.

### Task-evoked BOLD responses

Task-evoked BOLD responses were analyzed in FSL using the General Linear Model (GLM). The main blocks were modeled at the stimulus periodicity, and the voice instructions were modeled as 3-s events at the beginning and end of each block (see Fig. 2). A canonical double-gamma hemodynamic response function (HRF) was convolved with the timeseries of tactile stimulus blocks and voice events. Also, the motion parameter estimates of both of the simultaneously scanned heads were included as nuisance regressors for both heads individually; in other words, both subjects’ models had their own as well as the other subjects’ motion parameters as nuisance covariates. The other head’s motion estimates were included to gain resilience against motion-related field or signal fluctuations extending from one head to the location of the other. The analyses included the entire two-head volumes, allowing quantification and visualization of subject-specific and shared activation patterns across the dyad. In a complementary methodological approach, we used independent-component analysis with the GIFT toolbox (http://icatb.sourceforge.net/) on the joint dual-head EPI data, and we assessed the temporal profile of the top extracted components against the experimental stimulus model.

## Results

### Dual-coil performance

**Figure 3a-b** shows a representative subjects’ normalized data for T1 and EPI sequences. The temporal signal-to-noise ratio (tSNR) was compared between resting-state scans of the two subjects imaged simultaneously with the 2-person coil and the same subjects imaged alone with standard 32-channel head coil. **Fig. 3c** shows the mean tSNR in a sagittal plane of a representative dyad and in a roughly corresponding plane of one of the subjects of this dyad measured individually. The scales of the color bars are different by a factor of 1.5, corresponding to the theoretical scaling factor of SNR resulting from the differences in acquisition bandwidth (inversely proportional to the square-root of the bandwidth). As expected, the tSNRs of the two-person measurements were almost 50% lower those of the single-subject measurements, with most salient drop of signal in the frontal cortices.

**Figure 3.**
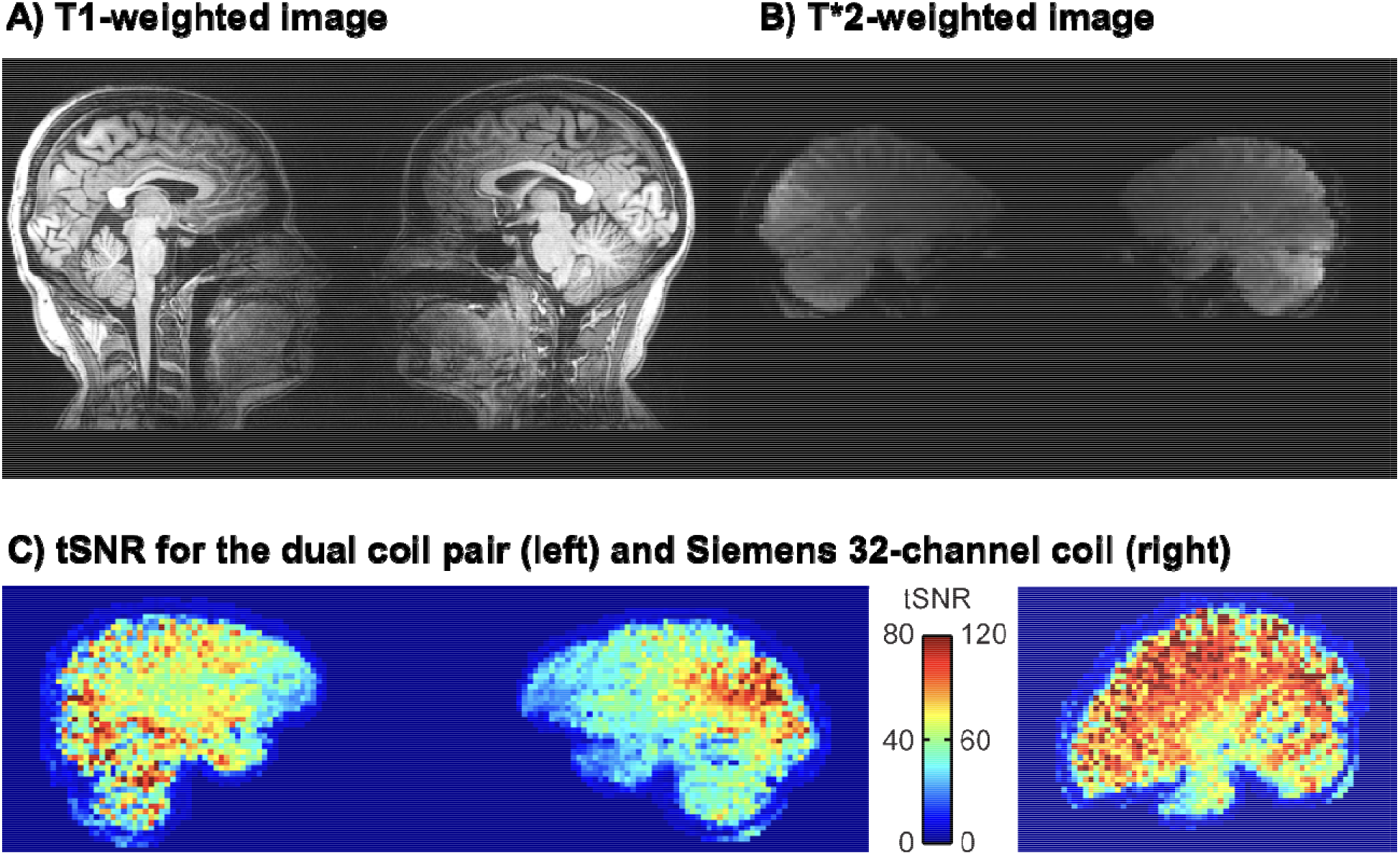
Representative single-pair T1 (A) and T*2 (B)-weighted images acquired with the dual coil. (C) tSNR for the dual coil and (D) conventional Siemens 32-channel head coil. Note that in due to preprocessing, the data from the dual coil pairs in panel (C) are further away from each other than they actually are (c.f. panel B)

### Regional Effects in the GLM

The voice cues modeled as 3-s events at the beginning of every rest and task command elicited reliable bilateral auditory-cortex activations similarly in both subjects regardless of their role as the actor or the receiver (**Fig. 4A**). In turn, the touching task resulted in differential activation patterns in the somatosensory and motor cortices depending on whether the subject was tapping or receiving taps (**Fig. 4B**).

**Figure 4.**
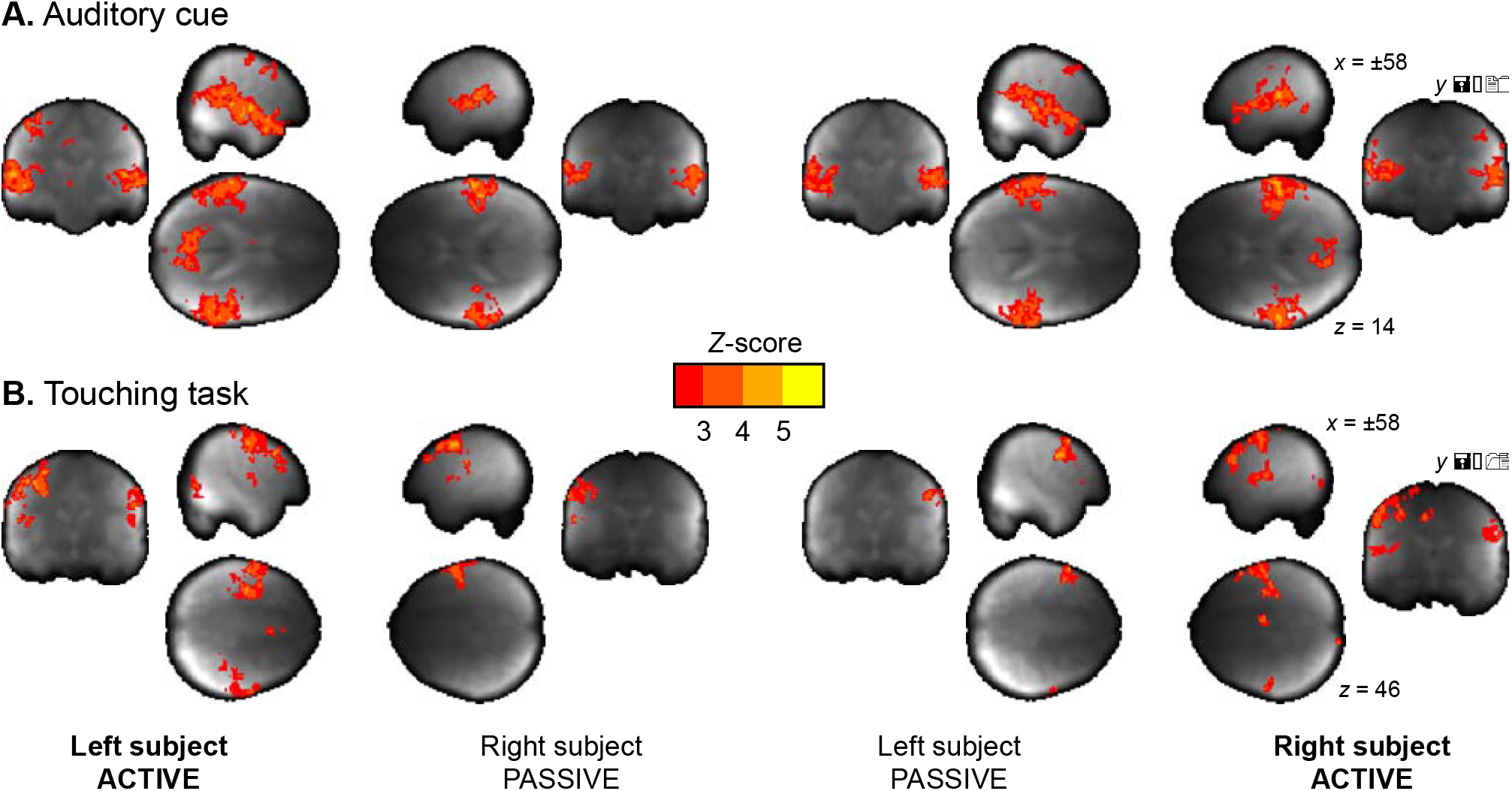
Main effects of verbal instructions (A) and the tapping task (B) for the actor and receiver subjects. The data are thresholded at p < 0.05, FDR corrected

We next evaluated the consistency of the auditory and somatosensory activations across individual subjects. To that end, we binarized the first-level activation maps for the verbal instructions and tactile tasks), and generated cumulative activation maps where voxels indicated in how many subjects task-dependent activations was detected at the a priori threshold (Fig. 5). This analysis confirmed that the evoked auditory responses could be detected practically in all the subjects, while the magnitude and detectability of the somatosensory responses was significantly more variable.

**Figure 5.**
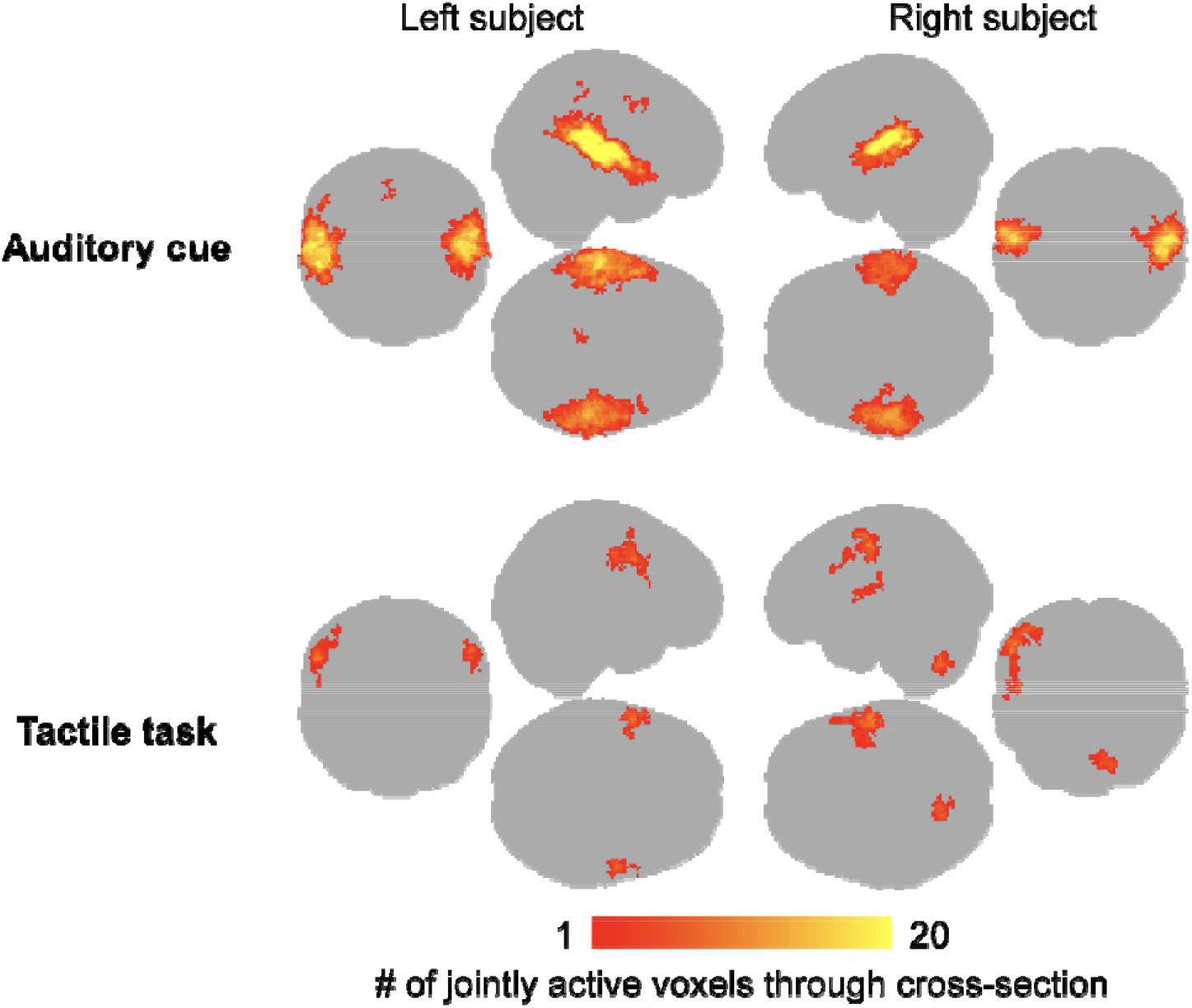
Cumulative map of the binarized (active / inactive) single-pair level activation maps for the auditory cues and tactile task. Colourbar indicates the number of subjects where significant activations were observed in the first-level analyses. Note that this analysis does not differentiate which subject was active in the tapping task.

### Independent-component analysis (ICA)

ICA (**Fig. 6**) applied on the combined data of the two subjects revealed two main components during the task: IC1 centrally involving the sensorimotor network, and the IC2 the auditory cortices and lateral frontal cortices. Component time courses tracked closely the stimulation model: The auditory IC2 had clear peaks at the onsets of the start and endpoints of each block, while the sensorimotor IC1 showed more sustained responses throughout the blocks. Both these components were shared with the subjects, implying that similar auditory and somatomotor activity patterns were present in both subjects, irrespectively of whether they were currently executing versus feeling the touches.

**Figure 5.**
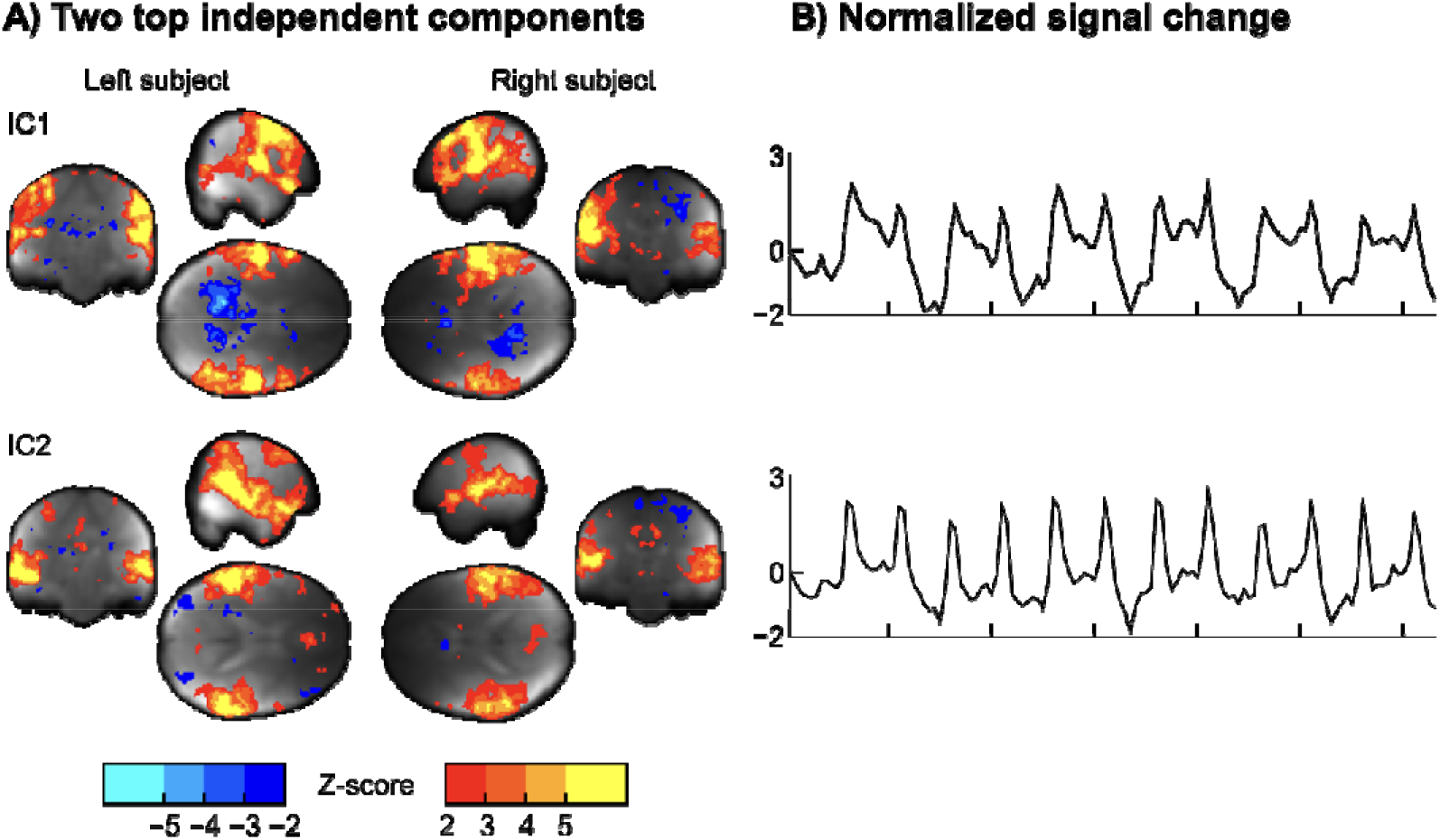
(A) Two top independent components (ICs) extracted from the data.

## Discussion

Our results show that haemodynamic activity can be reliably measured from two interacting subjects’ brains within one scanner using the dual-helmet setup, and that this technique can be used for studying elementary social cognitive functions, such as interpersonal communication via touching. Although the SNR of the dual-helmet coil was compromised (see Figure 3) compared with a conventional 32-channel head coil (Kaza et al., 2011), we could measure BOLD activations from both subjects. Task-dependent BOLD responses were task- and region-specific: auditory cues activated the auditory cortex similarly in both subjects (as they both heard the same cues), while the somatosensory and motor activations varied depending on which subject was actively tapping the other. The cues however appeared to alert the acting subject more than the reacting subject, as reflected by activation of the parieto-occipital cortex (precuneus). ICA also revealed similar task-dependent activation of sensorimotor and auditory networks in both subjects. Altogether our results highlight how sensorimotor networks ‘resonate’ across individuals during tactile interaction and confirm that the dual-coil fMRI is a potentially useful tool for studying brain basis of social interaction.

### Performance of the dual coil

Both GLM and ICA revealed that specific task-dependent fluctuations in haemodynamic activity can be picked up with the setup. However, SNR of the dual coil was clearly inferior to a conventional 32-channel head coil. An important source of discrepancy in the tSNR between the two- and the single-subject setups is the smaller number of coil elements in each of the helmets in the two-person coil in comparison to the one-person coil (8 vs. 32). The overall quality and geometry of the coil also matters: while the two-person coil is a working prototype, the 32-channel coil is the state-of-the-art product of the magnet vendor. The homogeneity of the main magnetic field (*B*_0_) is another important factor. The second-order shim coils cannot achieve the same degree of homogeneity for the two heads than for a single round object, and the *B*_0_ at the edges of the imaging volume is, to begin with, less homogeneous than in the centre of the magnet. For these reasons, the water peak is wider in the two-person case.

Also, as the two heads are typically of somewhat different size, the flip angles differ between the heads. Moreover, as the heads after shimming remain in different magnetic fields (and often result in a two-peaked water spectrum; the phase maps of the individual brains are relatively even, but have different offsets), the magnetization transfer due to fat saturation tends to reduce the signal of one head more than of the other, with fat saturation performance varying correspondingly. The homogeneity of the tSNR in the brain is also compromised due to the absence of coil elements in the anterior parts of the brains (see Figs. 1 and 3). This drop is similar to what occurs when the anterior part of the 32-channel coil is removed and only the posterior elements are used for imaging. A final reason that influences the tSNR is the subject comfort and stability, which in the two-person setup are worse than in the normal setup, still significantly compromised because the subjects need to be scanned in close proximity and in a sideways positions. We tried to alleviate this caveat by keeping the experimental runs short and by padding the subjects well, as well as using both subjects’ motion parameters as nuisance regressors on the analysis. It is however obvious that future studies need to implement more effective prospective means for motion control, such as neck or head restraints.

### Simultaneous measurement of interacting individuals

In contrast to conventional single-person MR imaging, the present two-person functional imaging approach provides novel means for understanding the neural basis of human social interaction. During social interaction, the interaction partners’ brains need to continuously anticipate, respond and adjust to incoming signals. A critical question is whether these sensorimotor loops function only recursively, as a cascade of third-person action-response processes? For example, a dialogue between two persons becomes fully incomprehensible if one persons’ speech fragments are removed from the recording. Brains are coupled with each other via behaviour, and influence each other via extracranial loops: Motor actions conveyed by one individual are interpreted by the sensory systems of another, and converted to sensorimotor format for promoting action understanding (Hari & Kujala, 2009). The present 2-P-fMRI setup provides means for studying how these loops are established during real-time interaction, as the evolving temporal cascade of sender-receiver operations in the social interaction can be measured continuously.

Intuitively the two-person neuroimaging sounds like the ultimate tool for analysing social interaction, because it allows quantifying the dynamic interaction between two brains similarly as such interaction occurs in real life. Yet after initial demonstrations of the feasibility of the two-person hyperscanning technique (Montague et al., 2002), it is surprising how little work has been conducted in this domain given the prominence of other individuals to practically all aspects of our lives (Dunbar, 1998). One likely reason is that that this type of studies are inherently complex to carry out and analyse. The two-person approach adds significantly to the complexity of the data – not just due to the doubled number of analysed voxels, but due to the interactive and temporally evolving nonlinear nature of real social interaction. It is thus possible that this line of work has not increased our knowledge on social interaction as much as the extra complexity would warrant. But it is also possible that we have not yet asked the best questions with the two-person neuroimaging setups, and maybe we need to adopt a new theoretical framework for measuring and analysing brain signals emerging from social interaction, rather than just scanning two brains at the same time using traditional approach with pre-determined stimulus models. During social interaction, the interlocutors constantly generate “stimuli” for each other in an adaptive fashion, so one potentially powerful approach involves careful recording and annotation of the behavioural dimensions of the social interaction as it occurs during the experiment, and using that data for post-experiment generation of the subject-specific stimulus models. This approach obviously leads to a high-dimensional stimulus space that again can be capitalized in the analysis: we do not necessarily know which features of social interacion form the most important dimensions when generating a classic stimulation model (Adolphs et al., 2016). On the contrary, when the stimulus model is generated based on the dimensions subject behaviour during the experiment, the critical dimensions do not need to be known in advance but the research may aim at constructing them based on the data.

### Practicality of the two-person imaging setup

We had to position our subjects into close proximity with each other due to the limited size of the transmitting body coil but also to provide a shared interpersonal space, allowing, for example, joint manipulation of objects. However, this intimate setting likely led to breaching the subjects’ peripersonal spaces, potentially influencing social processes because close social proximity with strangers may feel uncomfortable (Kennedy et al., 2009; Tsakiris, 2010). Accordingly, this setup is best suited for scanning subjects who know each other well enough, and the intimacy may also yield biases in subject selection. For the same reasons, this type of dual-coil imaging might also be difficult for patient populations with disorders involving social interaction. An optimized version of the coil design might simply involve a setup comparable to two conventional head coils arranged against each other on the dorsal end, so that both subjects can be scanned in supine position while they enter the scanner from opposite ends of the bore. Even though subjects cannot directly see each other, eye contact can be arranged using a mirror system. Our setup only had external auditory stimulus delivery system for the subjects. In theory, it would also be possible to project visual stimuli to the subjects, but due to the close proximity of the subjects’ faces this is deemed impractical. Our proof-of-concept study also revealed that the dual-coil setup is significantly less comfortable than conventional 32-channel head coil. Subject setup and shimming is slow, and the scanning position is difficult to maintain over prolonged periods of time. Because interlocking of the head coils and close proximity of subjects also increases susceptibility to motion. Accordingly, we tried to maximize subject comfort by limiting the scanning time into short blocks; in our experience the current scanning time (four 6-minute sessions plus anatomical images and preparations) was close to the maximum that subjects can comfortably do.

### Limitations and future directions

In this study we resorted to conventional moderately accelerated fMRI acquisitions. However, recent advances in multi-band excitation, to improve temporal resolution, and parallel transmit, to even out the flip angles in the two potentially very different sized heads, could greatly benefit the two-person MRI setup. The SNR for the dual coil was significantly worse than that of the conventional 32-channel head coil, particularly in the frontal cortex due to multiple factors pertaining to coil geometry and the low number of channels. This lacking signal in frontal cortex is a limiting factor when it comes to investigating social interaction, for which the frontal cortex acts as a central hub region (Amodio & Frith, 2006). However, many social processes emerge in regions where the coil system has adequate signal (such as posterior temporal and parietal cortex (Lahnakoski et al., 2012; Nummenmaa & Calder, 2009; Salmi et al., 2014), thus care must be taken when deciding what sort of social tasks can be studied with the present setup. Additionally, future benchmarking with variable tasks and experimental setups should be conducted to evaluate what types of tasks are ultimately feasible for this type of dual-coil imaging setup.

Future developments of the coil setup should strive to maximize coil coverage of the scalp more evenly, and with higher density coil arrangements. Such new devices would also allow more efficient control of subject motion: the limited contact of the current coil design with the scalp combined with the sideways scanning position makes the setup sensitive to head motion.

## Conclusions

We conclude that 2P-fMRI is a feasible and potentially powerful tool for studying brain dynamics of real-time social interaction. Even though the signal quality was compromised compared with state-of-the art head coils, our results show that it was sufficient for quantifying the dynamics of the real-time two-person interaction. This proof-of-concept study revealed that it is possible to measure good-quality haemodynamic signals simultaneously from two brains with one scanner. The two-person fMRI approach presented in this study complements the existing fMRI and MEG hyperscanning and face-to-face EEG and fNIRS techniques by allowing tomographic imaging of brain activations of two interacting subjects in face-to-face settings. Even though both subjects generated the tactile stimuli to each other in the experiment, the task was still externally controlled. Our data however suggest that in the future this methodology can be used for quantifying brain activation in dyadic, unconstrained, and naturalistic social interaction.

## Author Contributions

V.R., J.K., R.H., and L.N. designed research; V.R., S.M. and R.H. developed instruments, V.R. and J.K. acquired data, V.R. and J.K. analyzed data, V.R., J.K., R.H., and L.N. interpreted data, V.R., J.K., S.M., R.H., and L.N. wrote the paper.

## Acknowledgments

We thank Ms. Anna Anttalainen, Dr. Toni Auranen, Ms. Marita Kattelus, Mr. Veli-Matti Saarinen, and Mr. Tuomas Tolvanen for assististance, and Insight MRI for the development of the dual coil. The research was made possible by the Advanced Magnetic Imaging Centre, Aalto University School of Science, Espoo, Finland. V.R. is grateful to the funding provided by the Swedish Cultural Foundation in Finland, Instrumentarium Science Foundation, and Kalle and Dagmar Välimaa Fund of the Finnish Cultural Foundation. The funding support of the Academy of Finland (grant #218072 to R.H., grant #265917 to L.N.) and European Research Council (“Brain2Brain” grant #232946 to R.H. and “SocialBrain” grant #313000 to L.N.) is thankfully acknowledged.

---

**Supplementary Figure 1.**
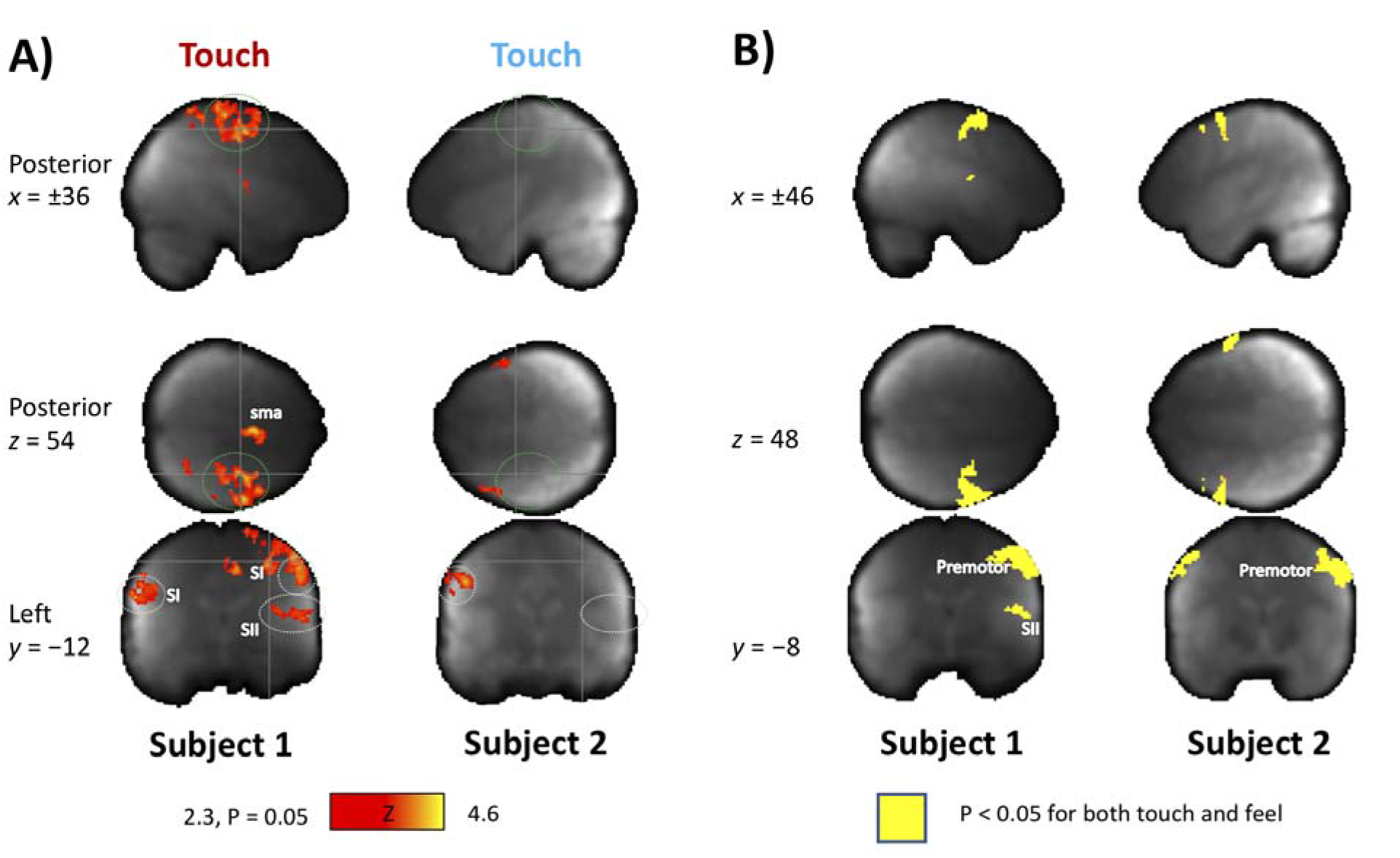
A) Alternating responses to active touching by left and right subjects. B) Overlapping activations for touching and feeling touch.

